# Uncovering the molecular basis of high morphinan product efficiency in opium poppy through Multi-omics integrated analysis with multi-capsules

**DOI:** 10.1101/2025.04.03.647041

**Authors:** Yumin Huang, Mao Sun, Wuhu Gong, Yifeng Liu, Yimeng Cheng, Lijuan Yuan, Yuanming Wu

## Abstract

Exploring the genetic landscape and corresponding regulatory mechanisms influencing morphine and related alkaloid production efficiency in the opium poppy (*Papaver Somniferum* L.) is important for sustainable medicinal production. However, information regarding the corresponding medicinal properties of poppy subspecies remains elusive, and the correlation between agronomic traits and morphine production remains unclear. Therefore, this study aims to investigate the distinction between two poppy subspecies: Shan Yang (SY) and Lan Tian (LT). Physiological data revealed that SY exhibited greater tolerance to environmental stress and a higher morphine content than LT, especially during the reproductive stage. To investigate genetic and molecular mechanisms underlying different tolerance traits and morphine synthesis efficiencies, a comprehensive comparison of genomic, transcriptomic, and metabolomic data was conducted, showing significantly higher single nucleotide polymorphism (SNP) frequency in SY (23.78%) compared to LT (23.69%). The proportions of SKIP (Skipped Exon) and MIR (Multi Intron Retention) were higher in SY than in LT. SY showed significantly higher gene expression related to benzylisoquinoline alkaloid synthesis, especially those involved in the steps from 1,2 dehydroreticulinium to morphine, narcotoline and noscapine, compared to LT. The upregulated expression of phenylpropanoid biosynthesis-related genes contributes to its more resistant agronomical traits. It plays a synergistic role in alkaloid production. The higher morphine content in SY resulted from an integrated effect controlled at different levels. Summarily, our findings offer new insights into the molecular mechanisms of morphine synthesis and present valuable gene resources for improving poppy cultivars with higher morphine content.

## 1. Introduction

The opium poppy (*Papaver Somniferum L*.) is a well-known medicinal plant with beneficial and harmful effects on human health [1]. Opioid-based analgesics, which are the main components of the opium poppy, remain among the most effective and inexpensive treatments for the relief of severe pain and palliative care. However, their potential for misuse due to their addictive properties poses a significant challenge. Opium-derived alkaloids, such as morphine, codeine, and thebaine, remain in great demand, especially in middle-to-low-income countries [2]. Therefore, high-yielding morphine and its related varieties are urgently needed to meet global demand.

Opium poppy remains the only commercial source [3] of any of the morphinan subclasses of benzylisoquinoline alkaloids (BIAs) because chemical synthesis or synthetic biology approaches have not attained commercial viability [4]. Various poppy subspecies have been identified worldwide [5, 6]. And environmental stress is crucial for opium poppy cultivation or the production of alkaloids. Typically, the natural mutant *Papaver somniferum* (*pap1*) exhibits significantly higher papaverine content than normal plants and follows different pathways for papaverine biosynthesis [7]. Poppy subspecies associated with domestication exhibit broad differentiation primarily based on capsule features, plant stature, and seed weight [8]. These physiological characteristics show diverse adaptations and responses to environmental stress. However, information regarding their corresponding medicinal properties remains elusive, and the correlation between agronomic traits and morphine production remains unclear.

BIA metabolism is a limiting factor in the production of morphine and related alkaloids. Massive genome expansion contributes to high BIA biosynthesis [6]. Furthermore, gene duplication, rearrangement, and fusion events are significant contributors to the evolution of BIA metabolism in the opium poppy. The P450 oxidoreductase gene fusion guides metabolites toward the morphinan branch and away from the noscapine branch [9, 10]. Essential for morphinan biosynthesis in the opium poppy, the combined P450 and oxidoreductase genes form the *STORR* gene fusion [1]. Furthermore, genetic modifications targeting the *COR* gene (codeine reductase) family cause the accumulation of the nonnarcotic alkaloid reticuline instead of morphine [11]. However, existing genome assemblies and comparative transcriptome analyses fall short of comprehensively capturing the genetic variations responsible for important agronomic traits, such as stress resistance and morphinan metabolism, within the subspecies. The relationship between the stress resistance of poppies and its contribution to BIA metabolism remains ambiguous.

Therefore, this study aims to investigate the distinction between two poppy subspecies (*P. somniferum*): one with a single capsule (Lan Tian: LT) and the other with multiple capsules (Shan Yang: SY) through a systematic multi-omics integrated analysis, including whole-genome sequencing, comparative transcriptome and metabolome analyses. The key variations responding to the agronomic traits related to environmental stress were identified not only across multiple organs and developmental stages in both subspecies, but also at different levels from genotype to phenotype.

## 2. Materials and Methods

### 2.1 Plant materials and physiological studies

Two Opium poppies were obtained from the Xi’an Public Security Bureau of Shaanxi Province. One was cultivated in Shanyang County, located in the southern region of Shaanxi province. This area boasts an average altitude of 1000 m, annual average sunshine duration of 2452.7 h, and annual average rainfall of 709 mm. It was designated SY. The second poppy was planted in Lantian county, situated in the northern part of Shaanxi province. This area has an average altitude of 1800 m, annual average sunshine duration of 2148.8 h, and annual average rainfall of 720.4 mm. It was labeled LT. All fresh samples used in this study were kept from the cases conducted in the Xi’an Public Security Bureau. The Xi’an Public Security Bureau approved all experimental procedures. Supplementary Table S1 shows details of these two subspecies. Agronomic traits, including plant height (from the root to the highest point of the plant), capsule number (representing the number of mature capsules), stem diameter (the diameter of the main stem), and capsule diameter (the maximum diameter of mature capsule), were measured.

### 2.2 Alkaloid content determination

Capsules were selected for alkaloid extraction, and the various alkaloids were isolated using a gas chromatography-mass spectrometer (GC-MS) (5975C-7890A, Agilent Technologies, USA) according to standard procedures. The alkaloid content of each sample was determined using at least three individual samples, and the data were expressed as mean±SD. Differences were assessed using the least significant difference (LSD) test, with statistical significance set at P<0.05.

### 2.3 DNA extraction, library construction, and genome sequencing

High-quality genomic DNA was extracted from freshly harvested leaves using the CTAB method, and the quantity and quality of the DNA were assessed using spectrophotometric analysis (NanoDrop 2000, Thermo Scientific, USA). Three types of short-insert paired-end libraries were constructed using the NEBNext Ultra DNA Library Prep Kit (Illumina, San Diego, CA, USA), with insert sizes of 200-, 300-, and 450 bp. Four types of mate-pair libraries, including 2, 5, 10, and 20 kb libraries, were constructed using Illumina DNA library preparation kits, following the manufacturer’s instructions. Genome sequencing was performed on an Illumina HiSeq platform (Illumina, San Diego, CA, USA).

### 2.4 Genome assembly and mapping to the reference genome

Raw data were filtered using SOAPnuke (V 1.6.5) to eliminate low-quality adapter contamination and PCR duplication. Supplementary Table S2 shows the resulting clean reads. The clean reads were then analyzed using Kemerfreq to generate the K-mer for subsequent analysis, employing a K-mer size ranging from 19–31bp. The Jellyfish software (V2.1.4) (http://www.genome.umd.edu/jellyfish.html) was used for rapid K-mer frequency count. GenomeScope software was employed to fit the k-mer spectrum to estimate genome characteristics. Fig. S1 shows the results. Poppy genome sizes were assessed using Genomescope (2.0) (http://qb.cshl.edu/genomescope/) software. Quantification was assessed using genetic maps of the opium poppy (HN1) published using ALLMAPS. The assembly quantity was measured by mapping the Illumina reads to the reference genome (HN1) (Table S3).

### 2.5 Phylogenetic analyses

To investigate the evolutionary history of SY and LT, opium poppy subspecies (HN1 and CHM) with published genome data were selected as references using OrthoFinder with default parameters, generating a matrix for phylogenetic analysis. Single-copy orthologs were identified from this dataset and utilized to construct a polygenetic tree.

### 2.6 Interspecific SNP effect comparisons

Single nucleotide polymorphisms (SNPs) were identified using GATK (version 3.8.1), leveraging alignments of short reads from HN1 onto the assembled SY and LT genomes. Subsequently, we identified heterozygous SNP sites in the SY and LT genomes based on the one-to-one genome alignment results. SNPs were filtered based on the following parameters: call quality divided by depth (QD), 2.0; mapping quality (MQ), 40.0; Fisher’s exact test (FS), 60.0; minor allele frequency (MAF), 0.05; and missing genotype rate, 0.2. Supplementary Fig. S2 and Table S8 show the results.

### 2.7 Methods for the bottleneck analysis and positive selection scan of the coding sequence

We utilized customized HBL wrappers for the TestBranchDNDS.bf HyPhy method files to assess evidence of the bottleneck effect on genome-wide Ka/Ks (the ratio of nonsynonymous to synonymous substitution rate). A scan of positive selection was conducted on the genes using the Ka/Ks ratio. The nonsynonymous to synonymous substitution ratio (Ka/Ks) serves as a metric for assessing the magnitude and direction of selection on coding sequences.

### 2.8 Characterizing patterns of selection and drift on gene expression

SY, LY, and two previously published poppies (HN1 and CHM) were employed for phylogenetic continuous trait modeling of gene expression. The median chronogram resulting from Bayesian tree reconstruction was utilized for this analysis. The model-fitted gene expression values were set log2-transformed before the analysis. We employed a model comparison approach, comparing Ornstein-Uhlenbeck (OU) and Brownian Motion models, to examine the influence of drift and different modes of selection on gene expression.

### 2.9 RNA extraction, library construction, and RNA sequencing

The whole plants were harvested in liquid N_2_ and stored at −80□. Frozen tissues were ground into a fine powder in liquid N_2_, and total RNA was extracted using TRIzol reagent (Invitrogen, USA), following the manufacturer’s instructions. RNA quantity and quality were analyzed through spectrophotometric analysis (NanoDrop 2000, Thermo Scientific, USA). Additionally, to ensure the integrity (RIN value≥8.0) of the RNA samples, analysis was conducted using the Agilent Bioanalyzer 2100 (Agilent Technologies) with the RNA 6000 Nano kit. Equal quantities of total RNA from three biological replicates were combined. Subsequently, they were used for cDNA library construction. The cDNA libraries were constructed using the Illumina TruSeq RNA Sample Prep Kit (Illumina Inc., San Diego, CA, USA) according to the manufacturer’s protocol. High-quality cDNA libraries suitable for sequencing were generated using an Illumina HiSeq 2500 sequencing platform (RiboBio Co. Ltd., Guangzhou, China). Furthermore, the raw sequencing reads were transformed into sequenced reads, or Raw Reads, through CASAVA Base Calling analysis. Subsequently, these reads were filtered again to obtain high-quality, clean reads. The quality metrics of the clean data, including Q20, Q30, and GC content, were evaluated (Table S9).

### 2.10 Functional annotation of the assembled transcripts

The Filtered Gene Set was identified by analyzing the alignment output files from TopHat. The resulting read counts were normalized using FPKM (fragments per kilobase of transcript per million mapped reads to quantify the transcript expression levels. Additionally, the R statistical package software edgeR (www.bioconductor.org/package/2.12/bioc/html/ed|geR.html) was used to identify high-quality differentially expressed genes (DEGs). DEGs were identified by comparing FDR (false discovery rate) value < 0.5 (http://www.bioconductor.org/packages/2.12/biochtml) and |logFC |≥1.

### 2.11 Gene region distribution statistics

The sequence obtained from sequencing was compared with the gene region where in the reference genome using the gene annotation files of the species available in the relevant database.

### 2.12 Alternative splicing analysis

The AStalavista software (https://confuence.sammeth.net/display/ASTA/2+-+download) was used to identify and categorize alternative splicing events in transcripts. Based on known gene models, alternative splicing is classified into the following categories: Transcription Start Site (TSS), Transcription Terminal Site (TTS), Skipped Exon (SKIP), Multi-exon SKIP (MSKIP), Intron Retention (IR), multi-IR (MIR), and Alternative Exon Ends (AE).

### 2.13 Relative expression of differentially expressed genes from enriched pathway analysis

RNA-seq reads of SY and LT were mapped to the opium poppy genome using TopHat (version 2.0.8), followed by KEGG pathway analysis to identify genes exhibiting different expression levels. Unigene expression was quantified using the FPKM method. DEGs were identified by applying the criteria FDR ≤ 0.001 and an absolute value of log2Ratio ≥ 1 (Table S12).

### 2.14 BIA and Phenylpropanoid biosynthesis pathway analyses

BIA-related and phenylpropanoid biosynthesis-related gene sequences obtained from the NCBI database (https://www.ncbi.nlm.nih.gov) were used as target sequences. These genes are known to be involved in BIA pathways, and the phenylpropanoid biosynthesis pathway has been previously cloned from HN1 and other plants, with experimental validation *in vitro*. We utilized these genes as search queries against SY and LT sequences using the BlastP algorithm, with an e-value cutoff of ≤1E−10. Only blast hits with >50% identity, and ≥80% coverage were retained and concatenated by solar. Conserved domains were identified in the retained sequences (Table S13 and S14).

### 2.15 Metabolite profiling

Seedlings cultivated by public security agencies were also collected. Secondary metabolites were identified. Following that, they were quantified using a hybrid quadrupole-TOF LC-MS/MS mass spectrometer (AB SCIEX Instruments, Shimadzu LC30AD). Each cultivar was evaluated with six biological replicates. Analysis was performed using an HQ-TOF LC/MS/MS system (Shimadzu, Japan), with detection using electrospray ionization (ESI) in positive and negative ion modes. Upon obtaining the chromatograms and mass spectrograms of the poppy compounds, qualitative analysis was conducted to assess the data using MS-DIAL 4.10 (Nature Methods, 2015). The identified compounds were cross-referenced against the published MassBank, Respect, and GNPS libraries. Principal component analysis (PCA) and partial least squares-discriminant analysis (PLS-DA) were conducted using SIMCAL14.1 (Umetrics 1987). Differentially expressed metabolites were screened by setting the p-value to ≤ 0.05 and fold change to (FC) ≥ 2 or ≤0.5, with VIP score ≥ 1.

### 2.16 RT-qPCR

DEGs were randomly selected from upregulated and downregulated genes. RNA samples for transcriptome analysis were reverse transcribed using a HiScript II 1st Strand cDNA Synthesis kit (Vazyme Biotech, Nanjing, China) with oligo (dT23) primers. RT-qPCR experiments were performed on a LightCycler480 Real-Time PCR System using ChamQTM SYBR® qPCR Master Mix (Vazyme Biotech, Nanjing, China), following the manufacturer’s instructions. Supplementary Table S17 shows the gene-specific primers.

## 3 Results

### 3.1 Agronomic Traits and Morphine Content

Two subspecies of *P. somniferum* (Table S1) exhibiting distinct agronomic traits such as tiller number, capsule size, plant height, stem diameter, and leaf shape were collected (Fig. 1A). SY showed better adaptability than LT attributable to its significantly longer stem diameter, more tillers, and larger capsules (Fig. 1B). The average heights of SY and LT were 124.6 cm and 108.6 cm, respectively. The average stem diameters of SY and LT plants were 1.99 cm and 0.99 cm, respectively). SY exhibited multiple capsules, while LT had only one (Fig. 1A and B). The primary benzylisoquinoline alkaloids in the capsule husk, including morphine, codeine, noscapine, thebaine, and papaverine, were measured using GC-MS. The average morphine and codeine contents were significantly higher in SY than in LT. However, no significant differences were observed in thebaine, papaverine, and noscapine contents between SY and LT (Fig. 1C).

**Fig 1.**
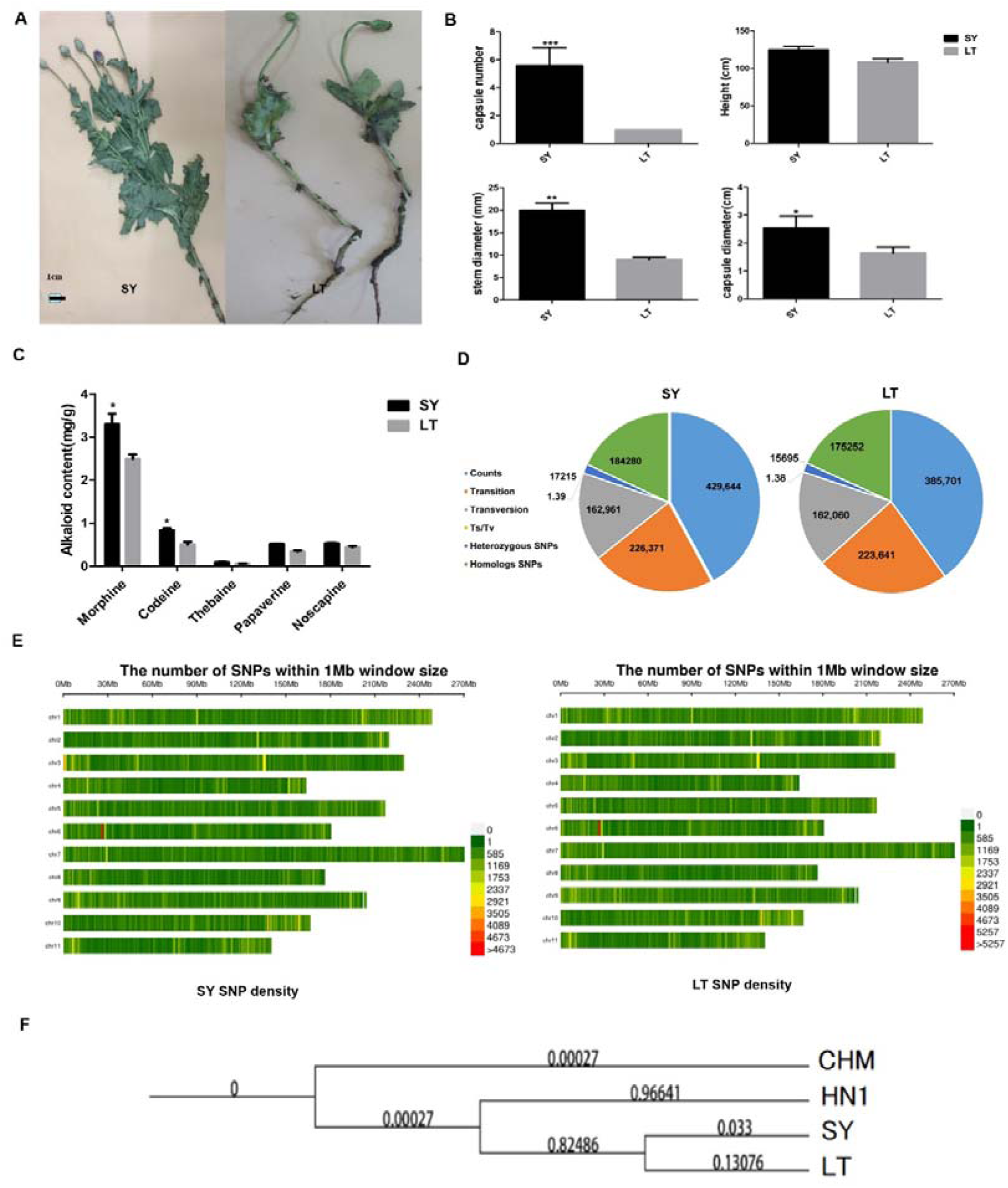
Comparative genomic analysis of multi-fruit cultivar SY and single-fruit cultivar LT. **(A)** SY and LT phenotypes. **(B)** The physiological characteristics of SY and LT. Values are presented as the means±SE (n=10). Ten plants were used to calculate the capsule number, plant height, stem diameter, and capsule diameter. Different letters indicate significant differences among treatments at P<0.05, according to the LSD test. **(C)** Alkaloid content of LT and SY. Different letters (a, b, c, d, e, and f) indicate significant differences (P < 0.05) according to the LSD test. **(D)** Results of SNP calling. **(E)** SNP density analysis results. **(F)** Phylogenetic tree of SY, LT, and two other published opium poppies.

### 3.2 Genome Assembly and Polymorphisms

To investigate the key molecular mechanisms affecting morphine production, SY and LT genomes were sequenced and assembled using a comprehensive strategy that involved DeNovoMAGIC-2, next-generation sequencing, and multiple mapping techniques. Approximately 162.27 Gb (∼61.52× genome coverage) of SY sequence data and above 173.19 Gb (∼65.66× genome coverage) of LT sequence data were obtained (Table S2). Using a negative binomial distribution model (Genoscope), the estimated genome size of LT was 2.90 Gb, with a heterozygosity rate of approximately 0.04%, and repetitive sequences accounted for approximately 65.59%. Simultaneously, the estimated genome size of SY was 2.92 Gb. The heterozygosity rate was approximately 0.04%, and repetitive sequences constituted approximately 65.73% (Fig S1). The quality of the assembled genome was evaluated using the HN1 genome as a reference. The mapping ratios for SY and LT were 96.13% and 96.71%, respectively (Table S3).

Genome polymorphisms generated in SY and LT were analyzed following a previously published method [12]. A total of 429,664 SNPs were identified in SY, comprising 226,371 transition sites and 162,961 transversion sites. Simultaneously, 385,701 SNPs were identified in the LT group, including 223,641 transition sites and 162,060 transversion sites (Fig. 1D and 1E; Table S4, Fig S2). The variant types and regions were further characterized. The Ts/Tv (Transition/Transversion) ratio in SY and LT was 1.39 and 1.38, respectively. A large proportion of downstream gene variants, non-coding transcript variants, and upstream gene variants were observed in the SY and LT genomes. Approximately 23.78% and 23.69% of the total SNPs were located in transcript regions for SY and LT, respectively (Table S4; Fig. 1D and 1E).

### 3.3 Phylogenetic Tree

A phylogenetic tree, which included SY, LT, and two other published opium poppy subspecies (HN1 and CHM), was constructed to clarify the evolution of the whole genome at the crucial nodes of *P. somniferum* divergence (Fig. 1F). The evolutionary pathways of SY and LT were more closely related to each other and showed a closer relationship to HN1 than CHM.

### 3.4 Genetic Variation and Positive Selection

To ascertain interspecific genetic variations between SY and LT, a series of experiments were conducted to identify disparities in gene expression and genome polymorphisms. Genetic distances between adjacent genes were calculated. SY and LT were treated as one group in the distance analysis because of their close evolutionary pathway, with HN1 serving as a reference control (Fig. 2a). The distance between adjacent genes exhibited considerable variation, indicating an increased probability of genetic mutation and linkage exchange between chromosomes in SY and LT during the evolutionary process (Fig. 2a; Table S5). The Ka/Ks substitution analysis showed a Ka/Ks ratio > 1, indicating that SY and LT were subjected to positive natural selection during evolution (Fig. 2B and C; Table S6).

**Fig 2.**
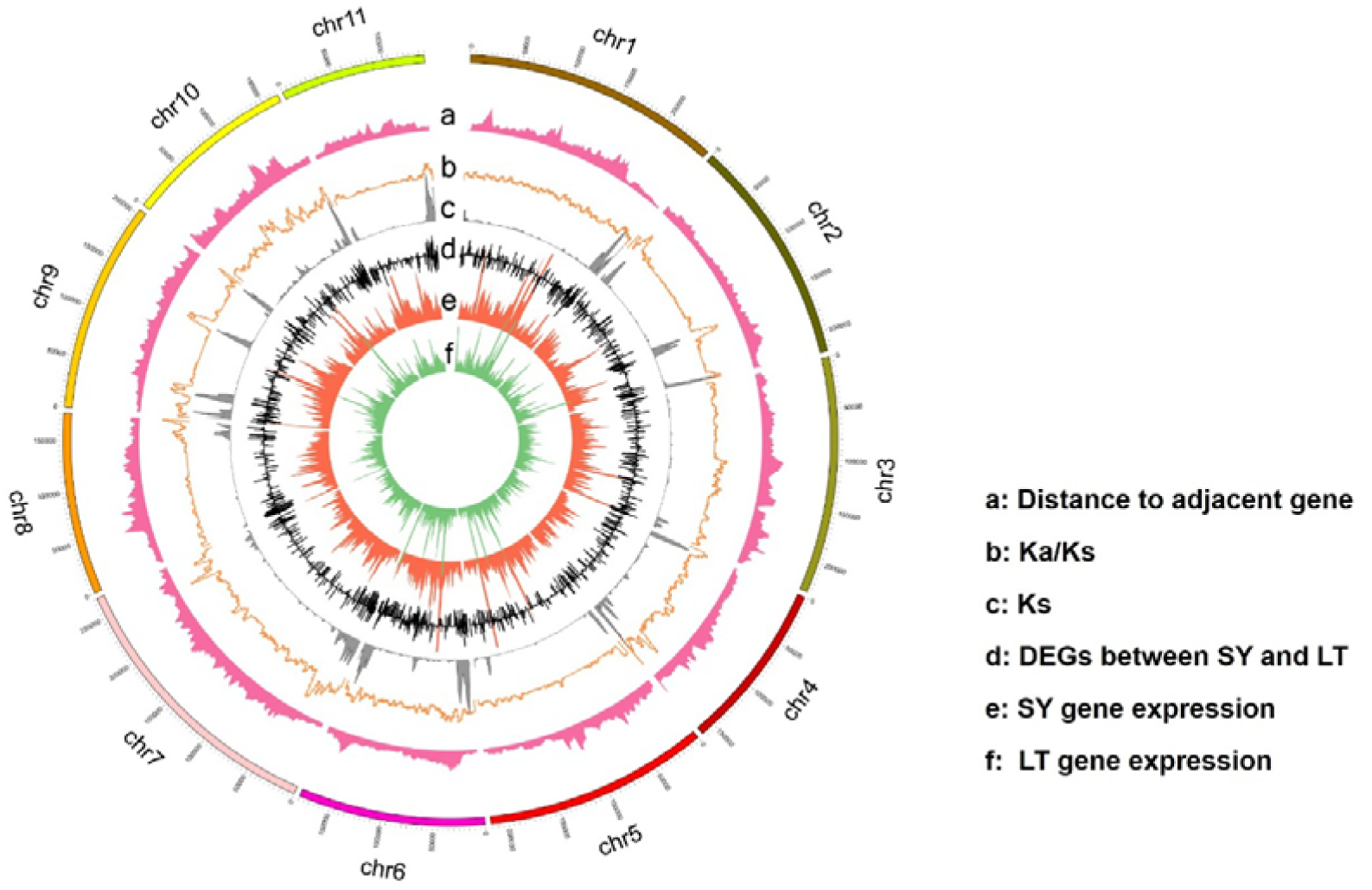
Diversity in SY and LT varieties. **(a)** Distribution of the mean distance to adjacent genes, with larger distances associated with centromeric sequences. **(b)** The ratio of nonsynonymous substitutions (Ka) to synonymous substitutions (Ks). **(c)** Synonymous substitution rate (Ks) distributions of syntenic blocks for SY and LT opium poppy species with *P. somniferum* paralogs. **(d)** Genes differentially expressed between SY and LT. **(e)-(f)** Gene expression of SY (e) and LT(f) samples.

A decrease in natural divergence (Ks) was observed near the centromeres, accompanied by a relative increase in Ka in the same regions. This pattern, as illustrated in the global patterns of nucleotide diversity (Fig. 2A–C; Table S6), suggests that SY and LT experienced active selection during the evolution process compared to HN1. Positive natural selection could contribute to the differences in secondary metabolites between SY and LT, specifically the higher morphine content in SY. Furthermore, the transcriptomes of SY and LT were analyzed according to a previous study, and 55,134 common transcripts were found in the two cultivars. Notably, 3,328 genes were expressed predominantly in SY, while 2,857 genes were expressed predominantly in LT. According to the opium poppy genome (HN1), genes located in centromere-proximal regions showed lower expression in SY than in LT, while the gene density in chromosomal arms was higher in SY than in LT (Fig. 2D–F; Table S7 and S8). This suggests that genes related to morphine synthesis in SY might be located on the chromosome arm.

### 3.5 Alternative Splicing Events

To further investigate whether the differential expression of genes on SY and LT chromosomes was related to morphine synthesis, gene fragments from transcriptome sequencing were aligned to the reference sequences (HN1). The mapped regions of SY were different from those of LT. Approximately 27,494,761 reads from SY, and 27,572,108 reads from LT were aligned to exon regions of HN1. In contrast, only approximately 495,279 reads from SY and 508,350 reads in LT were aligned to intron regions of HN1 (Fig. 3A). Approximately 12,000 AS events were identified in both SY and LT groups (Fig. 3C). The most abundant types of SY and LT were TSS and TTS (Fig. 3B, C). The combined proportion of TSS and TTS of total AS events in SY (35.75%+35.79%) was lower than that in LT (36%+36%). However, the proportions of SKIP and MIR were larger in SY (SKIP: 5.05% and MIR: 2.85%) than in LT (4.80% and 2.18%) (Fig. 3C). Interestingly, previous studies have indicated that during MIR and SKIP events, transcripts are not translated into protein. This phenomenon is believed to meet the needs for rapid plant growth through the rapid shear release in the later stages. The rapid changes associated with MIR can also serve as a mechanism for rapidly activating cell division and expanding fruit post-fertilization [13]. Therefore, it was hypothesized that the different frequencies of MIR events in SY and LT could influence the highly efficient production of morphine. This aligns with the findings suggesting that more MIR events not only accelerate plant cell division and growth but also promote capsule yield and effective secondary metabolite formation.

**Fig 3.**
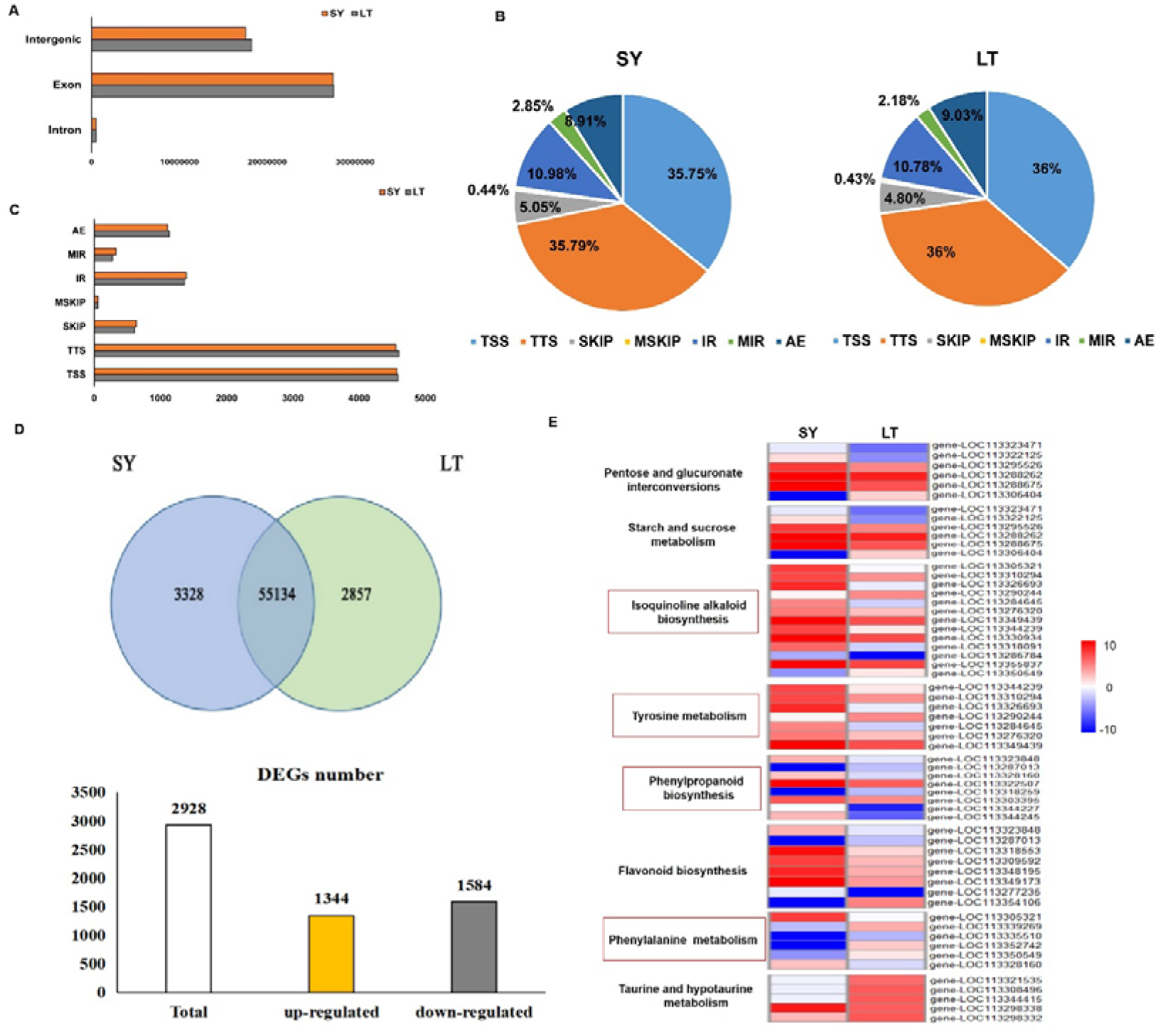
Comparative transcriptomic analysis of the molecular mechanisms of high morphinan product efficiency. **(A)** Distribution of SY and LT sequences according to the reference genome region. **(B)** Proportion of alternative splicing events in SY and LT. **(C)** Alternative splicing events in SY and LT. The x-axis represents the number of alternative splicing events, while the Y-axis indicates alternative splicing events. **(D)** Venn diagrams of DEGs in LT and SY and a summary of DEG numbers across both varieties. **(E)** Relative expression of differential genes in the enriched pathway analysis between SY and LT.

### 3.6 Gene Expression Events

To further investigate whether AS events led to more efficient production of morphine in SY than in LT, a comparative transcriptome analysis was conducted. The DEGs were identified based on FDR values < 0.05 and a log2 (fold change) ≥1. Overall, 1,344 genes were upregulated in SY, and 1,584 genes were upregulated in LT. (Fig 3D; Table S9–S11). KEGG analysis was performed using significant DEGs, and eight metabolic pathways were enriched, including pentose and glucuronate interconversions, starch and sucrose metabolism, isoquinoline alkaloid biosynthesis, tyrosine metabolism, phenylpropanoid biosynthesis, flavonoid biosynthesis, phenylalanine metabolism, and taurine and hypotaurine metabolism pathways (Fig. 3E; Table S12). The isoquinoline alkaloid biosynthesis, tyrosine metabolism pathway, and flavonoid biosynthesis pathway showed significant enrichment in SY. In contrast, the taurine and hypotaurine metabolism pathways exhibited significant expression in LT. Genes participating in isoquinoline alkaloid biosynthesis and tyrosine metabolism pathways were involved in morphine synthesis in the opium poppy, and the upregulation of these genes may contribute to the higher morphine content in SY.

Furthermore, the genes associated with the BIA pathway in SY and LT were analyzed. The expression of genes *NCS* and *4’OMT,* key enzymes in the synthesis of morphine and related alkaloid precursors, were upregulated in SY compared to LT. The genes of *SalAT*, *T6COM*, and *COR1,* involved in the morphine synthesis branch, exhibited upregulation in SY. This suggests the significance of the expression of these genes in morphine and related alkaloid synthesis. However, the expression of genes related to the noscapine synthesis branch showed inconsistencies. In SY, the expression of *CXE1, CYP82X2, and OMT* was upregulated compared to LT, while the expression of *CYP719A* and *CYP82X2* was downregulated (Fig 4; Table S13). This further confirms the crucial role of BIA-pathway-related gene expression in the higher morphine content observed in SY than in LT.

**Fig 4.**
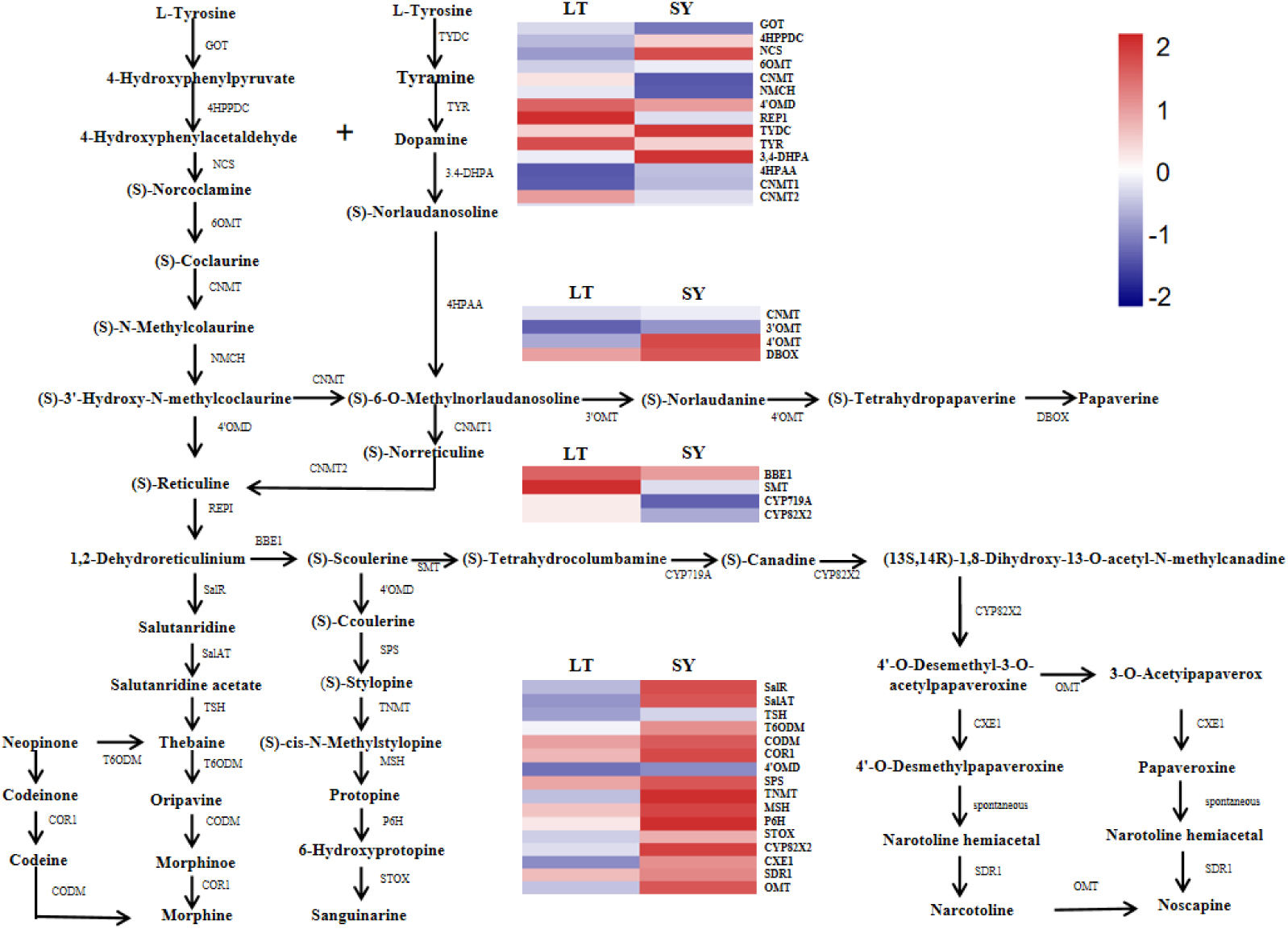
Gene expression in BIA metabolism pathway. Heatmaps illustrating gene expression levels in both t samples (SY and LT).

Phenylpropanoid metabolism has been reported to be related to plant growth and development, showing strong plasticity in response to constant environmental change [14, 15]. The KEGG analysis indicated an overrepresentation of phenylpropanoid biosynthesis in SY compared to LT. Among the 15 DEGs involved in this pathway, 11 exhibited upregulation in SY compared to LT. (Fig. 5; Table S14). Particularly, the genes *REF1*, *4CL*, *POX,* and *CCR1* play important roles in phenylpropanoid metabolism, serving important functions in plant resistance to biological and abiotic stresses [16–18]. Tyrosine is involved in both phenylpropanoid metabolism and the BIA pathway. To validate the RNA-Seq findings, ten DEGs with upregulated and downregulated expression were randomly selected for RT-qPCR verification. The results showed a consistent expression trend for all DEGs compared to the sequencing results, proving the reliability of the sequencing results (Table S17 and S18; Fig. S4).

**Fig 5.**
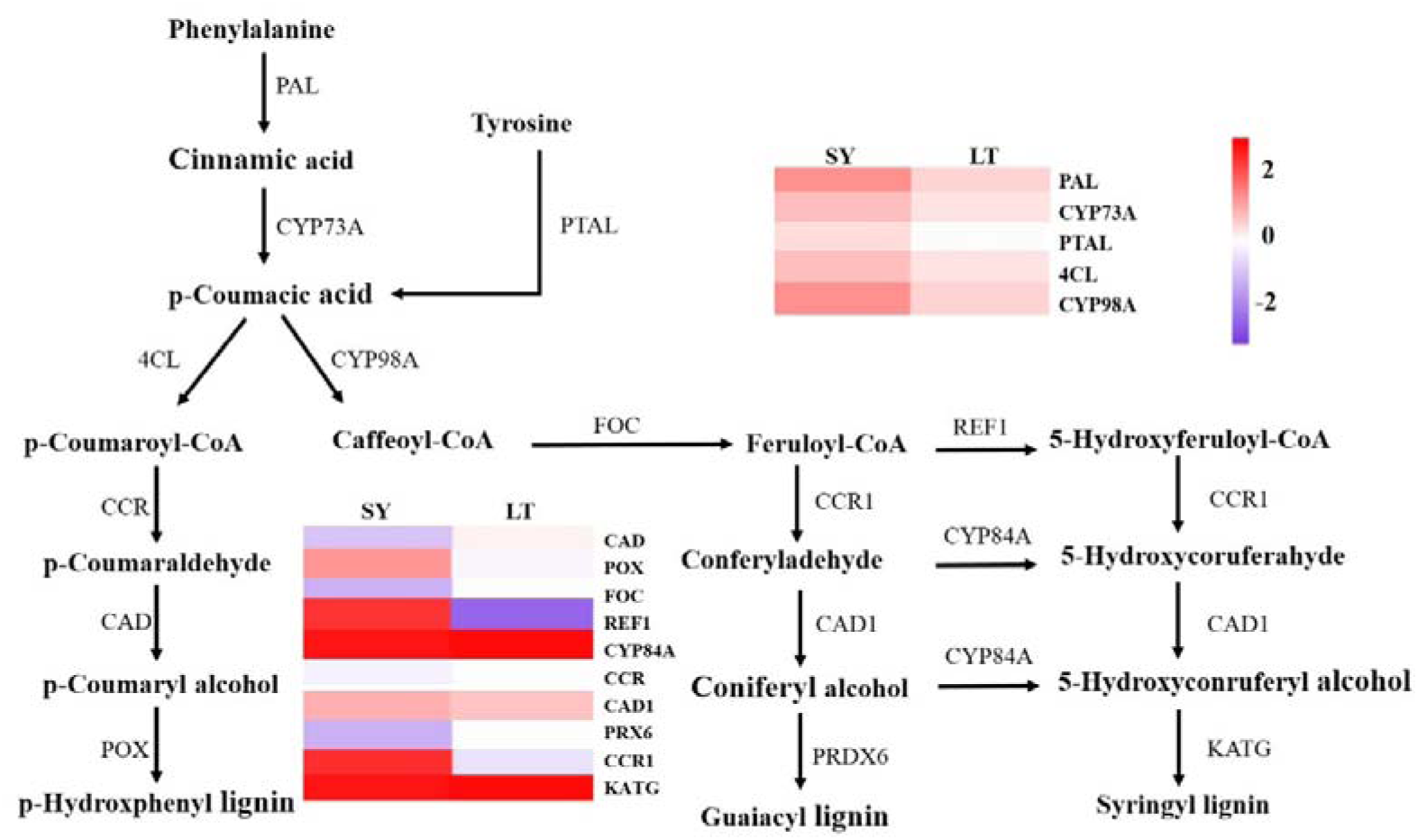
Gene expression in Phenylpropanoid metabolism pathway. Heatmaps showing gene expression levels in both samples (SY and LT).

### 3.7 Metabolism analysis Events

To better understand the differences in metabolites between SY and LT during growth and development stages, an analysis of the metabolite in the seedlings was conducted. The PLS-DA model revealed a distinct difference in metabolites between SY and LT (Fig. 6A). KEGG analysis revealed distinct enrichment of different metabolites in the flavone and flavonol biosynthesis and flavonoid biosynthesis pathways ((Fig. 6B, C; Table S15). The primary metabolites identified were located downstream of phenylalanine metabolism, as observed in the metabolite comparative analysis (Fig. 5 and 6D). Notably, SY exhibited higher levels of cinnamoyl-CoA, pinocembrin, and caffeoyl-CoA, while LT displayed upregulated levels of p-coumaroyl shikimic acid and naringenin (Fig. 6D; Table S16).

**Fig 6.**
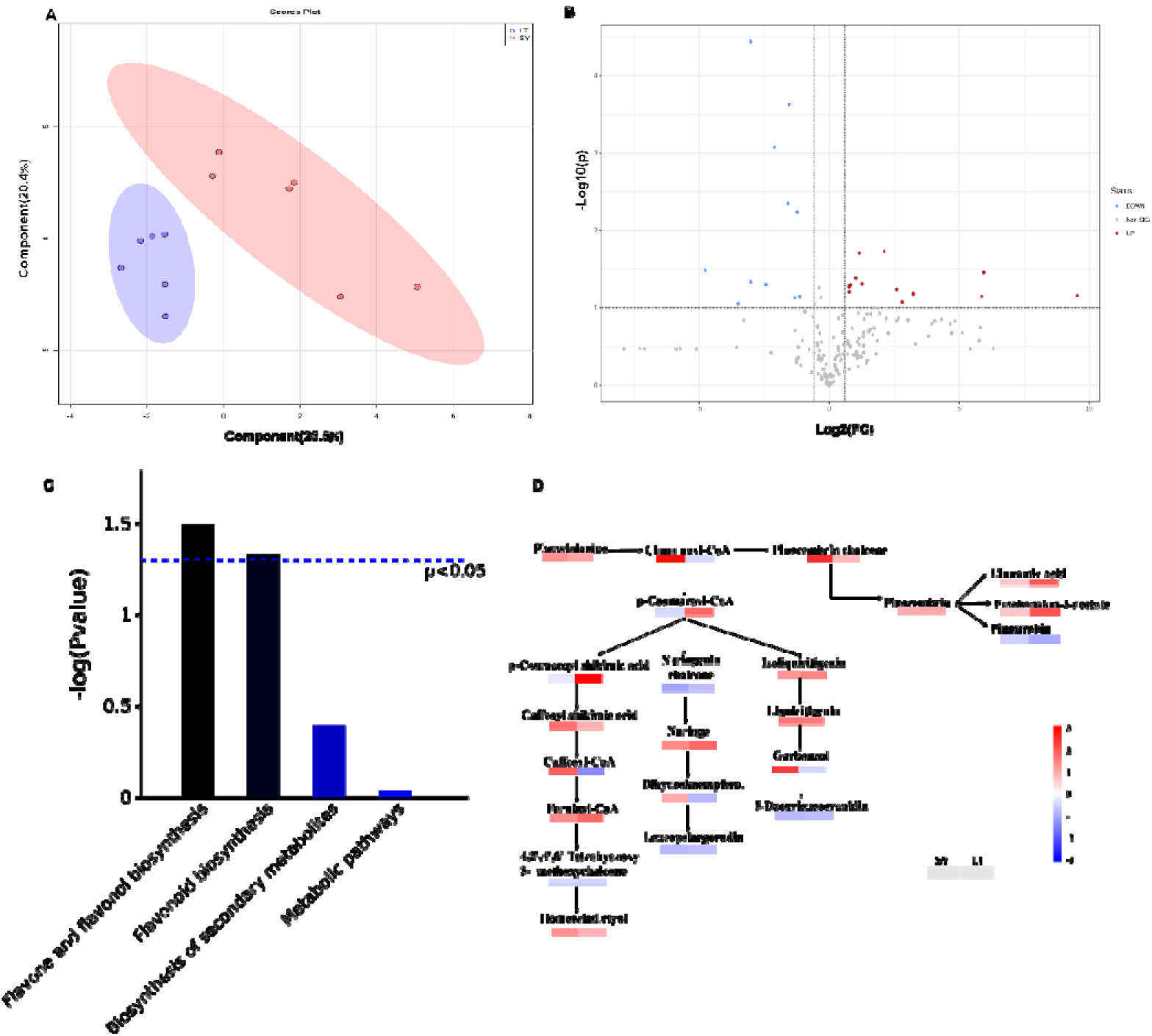
Comparative metabolic analysis of the seedlings of opium poppy. **(A)** PLS-DA score charts for SY and LT. The x-axis represents the score value of the main component, while the y-axis indicates the score value of the orthogonal component. The different points represent duplications of different samples. **(B)** Metabolism of SY and LT. The red and blue dots represent significant differences in metabolites. **(C)** Enriched KEGG pathways between SY and LT **(D)** Metabolism of the flavonoid biosynthesis pathway. Heatmap representing metabolism expression levels in two different samples (SY and LT).

## 4. Discussion

Opium stands out as the only source of morphine and related alkaloids, which globally significant for their pain-relieving roles. Various studies [1, 19] have explored various aspects, including genome comparisons, agronomically important traits, and the physiological and medicinal properties of poppies. However, the relationship between genetic properties and morphine content remains unclear. This study introduces two Chinese opium poppy landraces from Shaanxi Province, SY and LT, for the first time. Compared to LT, SY displayed heightened adaption to environmental stress, as evidenced by its strong stems, increased tillers, large capsules, and elevated plant height. Moreover, SY demonstrated a remarkably high morphine content. The distinct agronomic and medicinal differences between these subspecies shed light on the genetic factors contributing to high morphine content, offering valuable insights for enhancing *P. somniferum* germplasm, the most economically valuable crop for morphine production.

Limited research has been conducted on SNP analysis in opium poppies, and the gene localization of the identified SNPs remains unclear. The SNPs in SY and LT were predominantly concentrated in the downstream transcription region, with a higher mutation proportion in SY (23.78%) than that in LT (23.69%). This study marks the first report of SNP distribution in poppy subspecies with varying agronomic and medicinal properties, potentially providing genetic insights for the future traceability of opium poppies. This is pertinent for the timely detection of morphine-related cases and the eradication of illicit drug sources.

Furthermore, SY and LT demonstrated close evolutionary distances. Compared to HN1, the obvious variation in the distance between adjacent genes suggests an increased probability of genetic mutation and linkage exchange between chromosomes in SY and LT, probably increasing through the evolutionary process. The ratio of Ka to Ks (Ka/Ks) revealed a reduction in neutral divergence (Ks) near the centromeres but a relative increase in Ka in these regions, implying active selection through synonymous or non-synonymous mutations during natural evolution. Moreover, the proportion of SKIP and MIR in SY is greater than that in LT, suggesting adaptations for rapid plant growth through rapid shear release in later stages, and the particular way of evolution also shows up in functions. This process may explain the divergence of these two subspecies into distinct traits [20]. The contrasting genome landscapes of SY and LT enhance our understanding of poppy environmental adaptation and species protection. [21].

Gene expression patterns were also analyzed across different morphine and codeine concentrations in SY and LT. Pei et al. reported that the expression of genes in the noscapine biosynthesis cluster was lower in CHM than in HN1 [19]. In this study, KEGG analysis revealed a notable enrichment of pathways related to isoquinoline alkaloid biosynthesis and tyrosine metabolism in SY compared to LT. Key enzyme genes involved in the synthesis of 1,2-dehydroreticulinium into morphine and other important alkaloids (morphine, narcotoline, noscapine, and sanguinarine) showed significantly higher expression levels in SY than those in LT. These key enzymes, including *SalR*, *COR1*, *TNMT*, *CYP82X2*, *OMT*, et al., proved their crucial roles in the BIA pathway for morphine and related alkaloid synthesis [22, 23].

Especially, genes associated with phenylpropanoid biosynthesis exhibited significant enrichment in SY compared to LT. This pathway is related to plant growth and development with adaptability to constant environmental change [14, 15, 24]. Moreover, genes involved in downstream lignin synthesis, such as REF1, 4CL, POX, and CCR1, were significantly upregulated in SY compared to LT, facilitating water and nutrient transport [25]. Phenylpropanoid metabolites also enhance plant stress responses to abiotic and biotic stresses and serve to attract insects for pollination and seed dispersal [26, 27]. Dong et al. found that the expression of the PAL gene involved in phenylpropane metabolism in R rose varieties was significantly upregulated when aphids infected[28], a finding echoed in our study, where PAL expression in SY was significantly higher than that in LT, indicating heightened stress resistance in SY.

Metabolome analysis underscored the importance of phenylpropanoid metabolism, with flavonoid metabolism showing enrichment in seedlings from SY and LT, derived from phenylalanine through the phenylpropanoid pathway [29]. Therefore, we hypothesize that DEGs involved in phenylpropanoid biosynthesis may contribute to the distinct agronomic traits and medicinal properties of SY and LT, potentially interacting with the BIA pathways. Other metabolic pathways, such as pentose and glucuronate interconversion, starch and sucrose metabolism, flavonoid biosynthesis, and taurine and hypotaurine metabolism, also exhibited differential enrichment between SY and LT. These findings expand the metabolic database of the opium poppy and enhance our understanding of the differences between SY and LT, though further research is necessary to elucidate the specific roles of these pathways.

## 5. Conclusion

In conclusion, two poppy Chinese opium landraces from Shaanxi Province were reported for the first time in this study. The first comprehensive analysis of their agronomic and medicinal traits, alongside a detailed examination of their genomic, transcriptomic, and metabolomic features, were clarified using integrated analysis were provided. The more resistant agronomical traits of poppy (SY) may be attributed to genetic polymorphisms shaped during the evolutionary process and positive natural selection. These genetic changes may have also increased the synthesis of effective alkaloids, particularly morphine. This detailed genetic analysis not only establishes a foundation for future evolutionary and molecular studies of poppies but also holds significance for breeding programs and the enhancement of valuable alkaloid synthesis. Moreover, it further reinforced the pivotal role of genes in the BIA pathways for morphine and related alkaloid production. The upregulated expression of phenylpropanoid biosynthesis-related genes not only contributes to the more resistant agronomic traits but also synergistically influences alkaloid production (Fig 7).

**Fig. 7.**
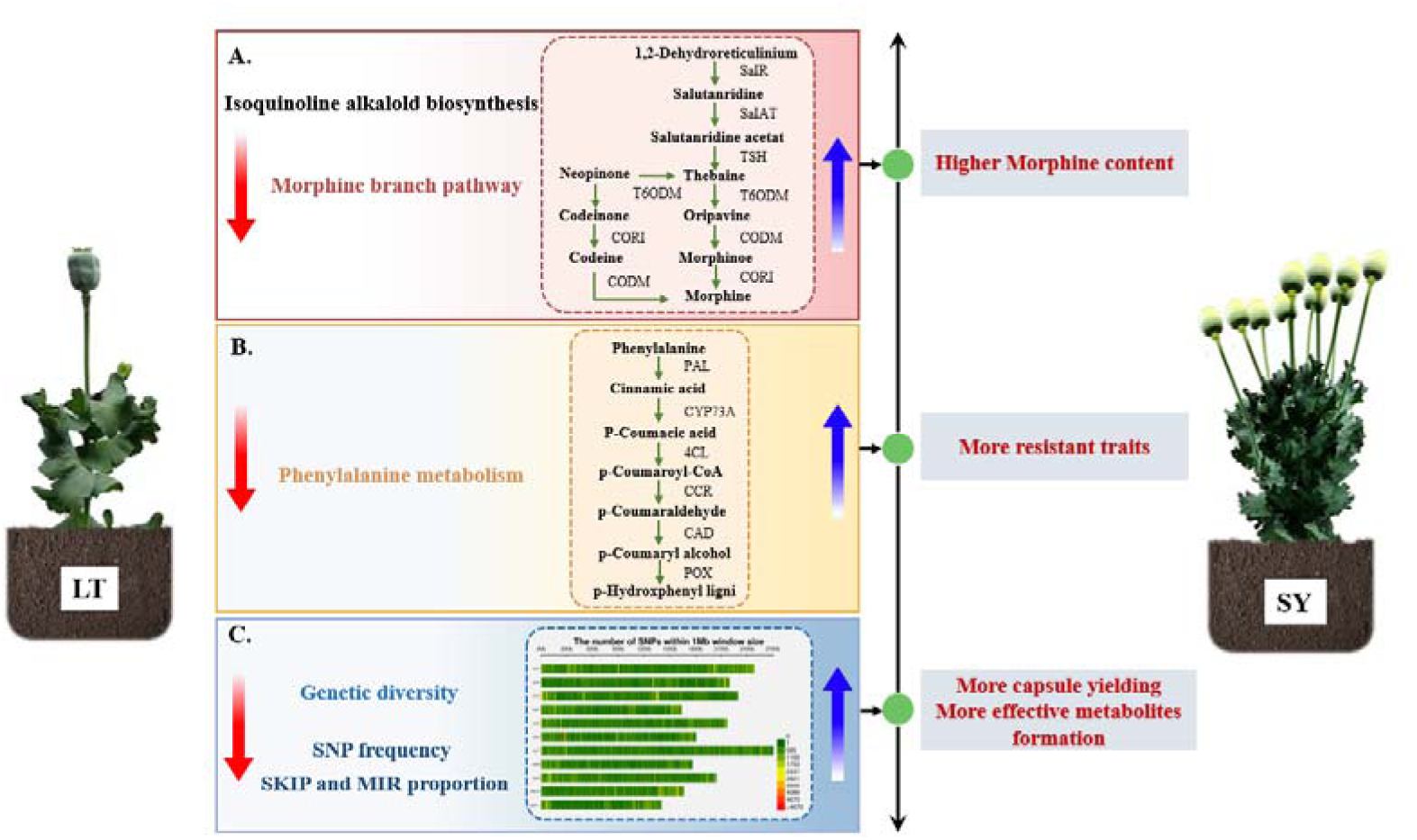
Schematic illustration of the main processes involved in differential morphine production in SY and LT plants.

## Abbreviations

BIAs: benzylisoquinoline alkaloids
pap1: Papaver somniferum
COR: codeine reductase
GC-MS: gas chromatography-mass spectrometer
LSD: least significant difference
SNPs: Single nucleotide polymorphisms
MAF: minor allele frequency
Ka: nonsynonymous substitution
Ks: synonymous substitution
OU: Ornstein-Uhlenbeck
TSS: Transcription Start Site
TTS: Transcription Terminal Site
SKIP: Skipped Exon
MSKIP: Multi-exon SKIP
IR: Intron Retention
DEGs: differentially expressed genes
MIR: Multi-IR
AE: Alternative Exon Ends
FPKM: fragments per kilobase of transcript per million mapped reads
ESI: electrospray ionization
PCA: Principal Component Analysis
PLS-DA: Partial Least Squares-Discriminant Analysis
AS: alternative splicing
LT: Lan Tian
SY: Shan Yang

## Acknowledgments

We appreciate Xi’an Public Security Bureau of Shaanxi Province for kindly providing plant materials. We thank Editage for providing English language editing of the manuscript.

## Sequencing data upload to NCBI

Accession number for sequencing data uploaded to NCBI: PRJNA994321.

## Author Contributions

The research concept was developed by Yuanming Wu and Lijuan Yuan. Yuming Huang conducted the bioinformatic analysis, Mao Sun contributed to the evolutionary analysis, and Wuhu Gong and Yimeng Cheng prepared the sequencing samples. Yifeng Liu participated in metabolite profiling and RNA-seq. The manuscript was drafted by Yuming Huang with the assistance of Mao Sun. All authors have thoroughly reviewed and approved the final version of the manuscript.

## Supplementary Information Appendix

### Funding and Acknowledgement

This work was supported by the key research and development plan in Shaanxi (Grant/Award Numbers: 2019SF-059 and 2020SF-204), the key innovative project in Shaanxi (Grant/Award Number: 2021ZDLSF02-02), and the National Natural Science Foundation of China (Grant/Award Number: 81671476). Additionally, the work received partial support from The Criminal Investigation Bureau of Xi’an Public Security Bureau.

### Disclosures

The authors have no conflicts of interest to declare.

### Supporting information

**Fig. S1.**
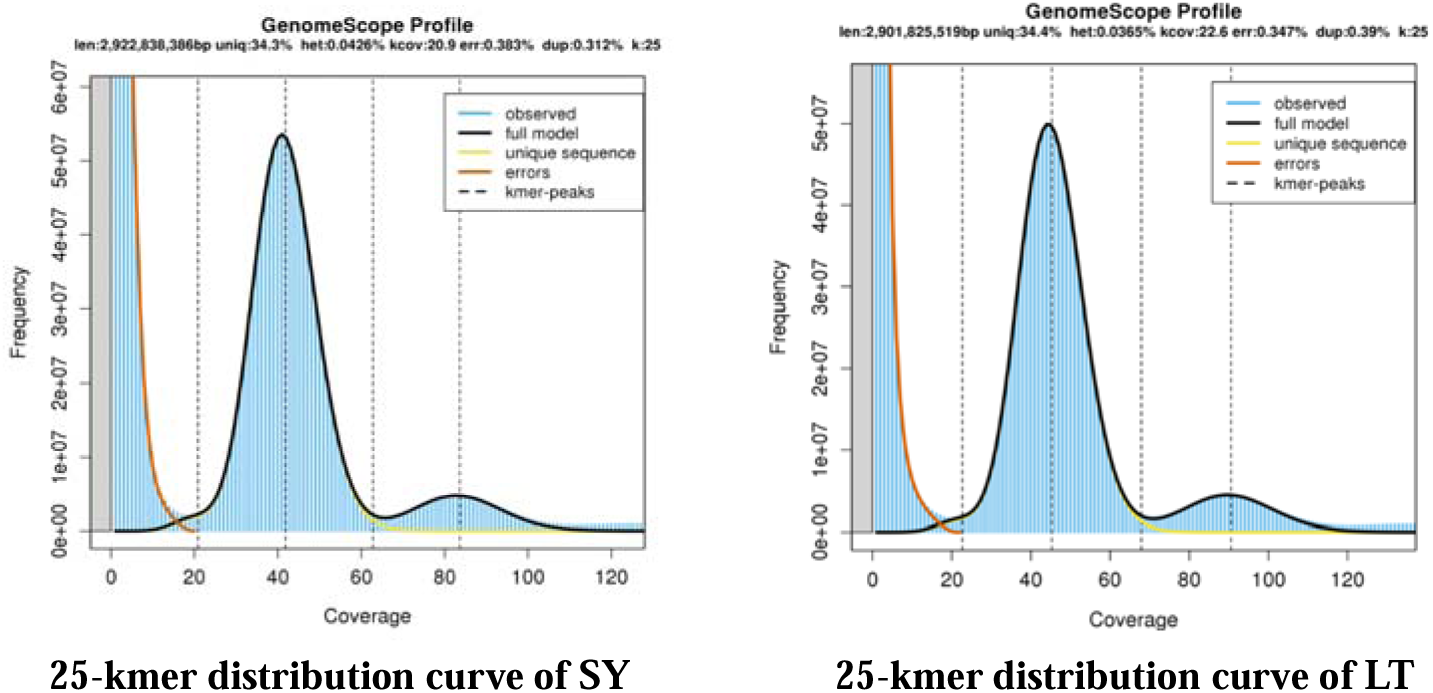
The k-mer depth distribution analysis of two cultivars. For better to estimate the characteristics of the genome,19-31bp multiple k-mer were analyzed, then, 25bp k-mer analysis was found with small sequencing bias and low error rate. The abscissa is the 25-kmer depth (coverage), and the ordinate is the 25-kmer frequency at this depth.

**Fig. S2.**
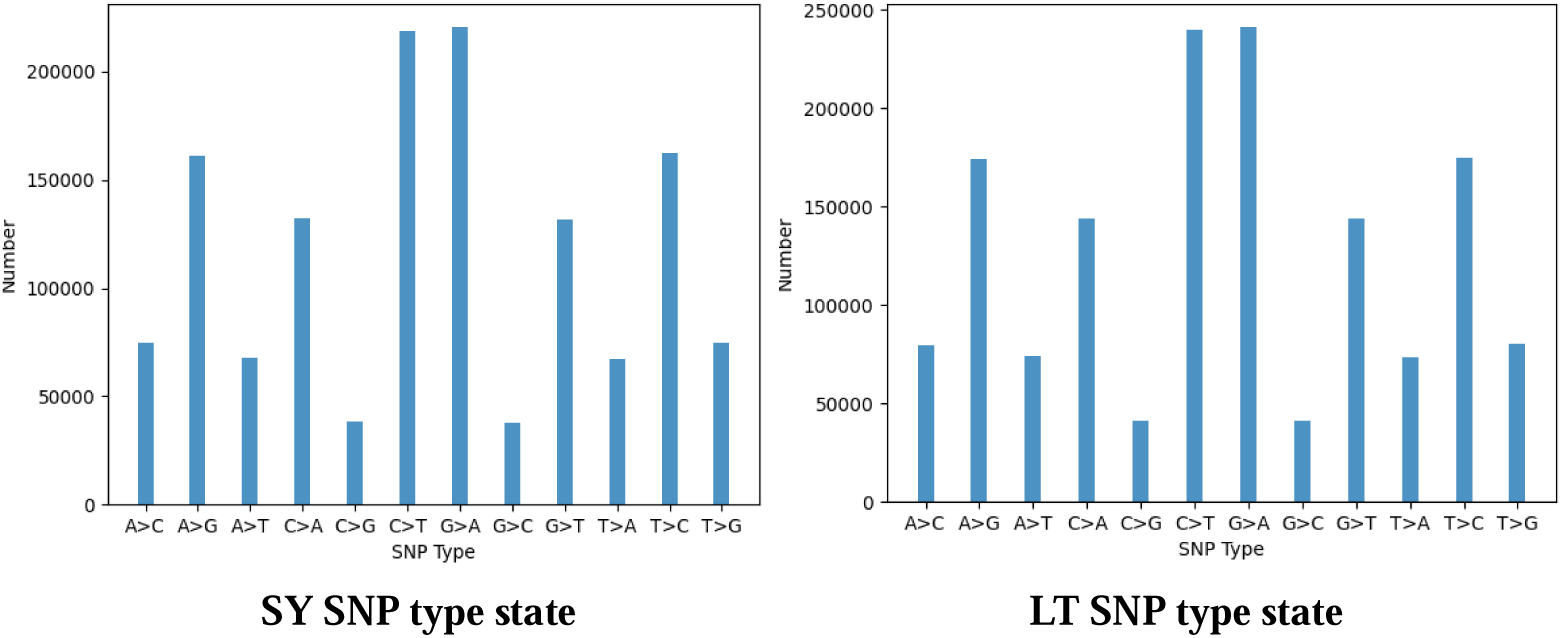
The result of calling SNP. Distribution of SNP polymorphisms along each opium poppy chromosome, data were generated via re-mapping the Illumina reads to opium poppy genome assembly, SNP substitution types in poppy genome. Y-axis gave the count of SNPs while the X-axis showed the indel length in nucleotides, for the LT species and SY species, respectively.

**Fig. S3.**
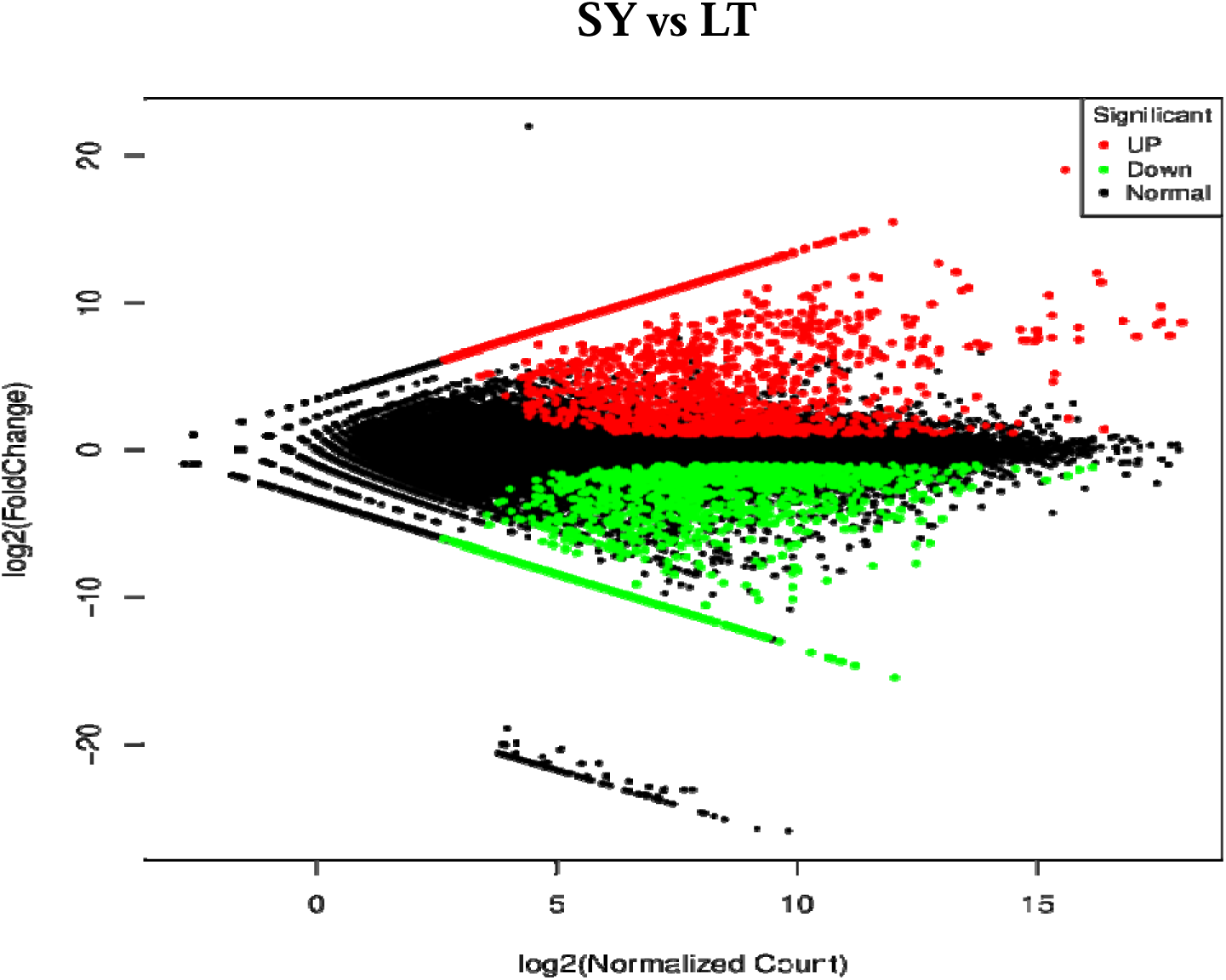
Distribution of DEGs between single-fruited(LT) and multi-fruited(SY) varieties. The volcano map of DEGs in the two varieties, red spots represent up-regulated DEGs and green spots indicate down-regulated DEG. The detailed information was shown in Table S11. (http://www.biocondutor.org/packages/2.12/biochtml/edgeR.html).

**Fig. S4.**
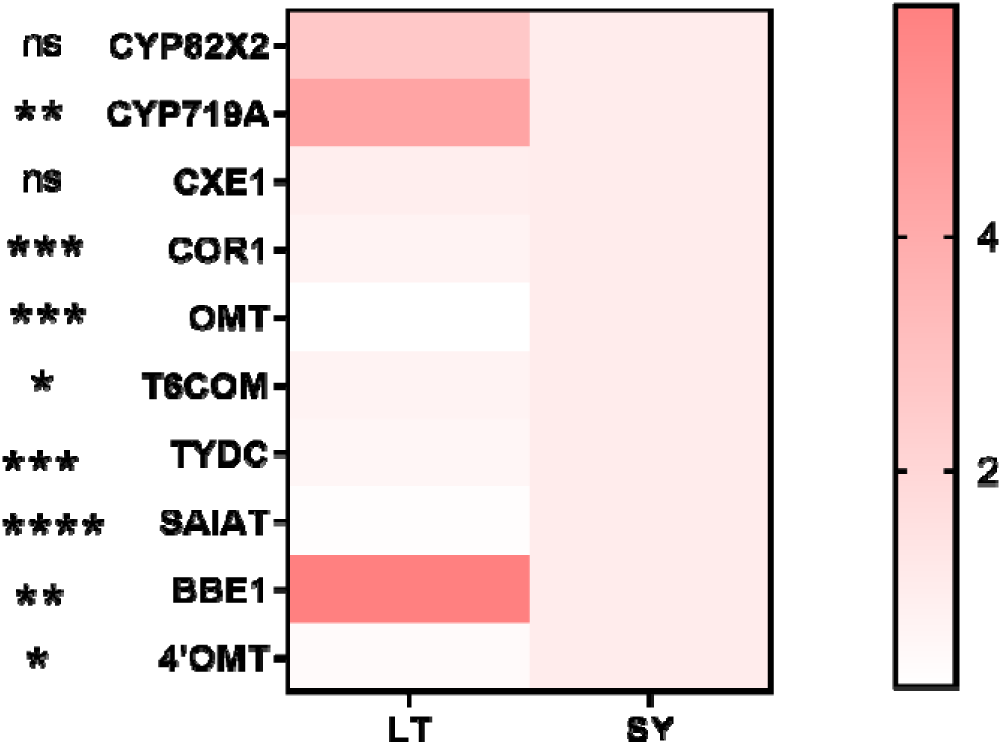
The heatmap of RT-qPCR results. Taking SY sample as control, LT as sample, the qPCR value of LT/SY indicates that DEGs in LT were up-regulated or down-regulated compared with SY. The detailed information was shown in Table S17, S18**.s**

Table S1. Material information.

Table S2. Filtered data statistics.

Table S3. Mapping of DAN-seq reads to the reference.

Table S4. SNPs count upon mapping to poppy genome..

Table S5. The gene distance.

Table S6. The ratio between Ka and Ks.

Table S7. The gene expression of LT.

Table S8. The gene expression of SY.

Table S9. RNA Sequencing output statistics.

Table S10. Mapping of RNA-seq reads to reference.

Table S11. The DEGs between LT and SY.

Table S12. The expression of DEGs in pathways.

Table S13. The gene expression of BIA between SY and LT.

Table S14. The gene expression of Phenylpropanoid pathway.

Table S15. The metabolites in SY and LT seedlings.

Table S16. The metabolites of Flavonoid pathway.

Table S17. The primers of DEGs of RT-qPCR.

Table S18. The results of RT-qPCR.

## Reference

[1] L. Guo, T. Winzer, X. Yang, Y. Li, Z. Ning, Z. He, R. Teodor, Y. Lu, T.A. Bowser, I.A. Graham, K. Ye, The opium poppy genome and morphinan production, Science, 362 (2018) 343–347.

[2] F.M. Knaul, P.E. Farmer, E.L. Krakauer, L. De Lima, A. Bhadelia, X. Jiang Kwete, H. Arreola-Ornelas, O. Gomez-Dantes, N.M. Rodriguez, G.A.O. Alleyne, S.R. Connor, D.J. Hunter, D. Lohman, L. Radbruch, M. Del Rocio Saenz Madrigal, R. Atun, K.M. Foley, J. Frenk, D.T. Jamison, M.R. Rajagopal, C. Lancet Commission on Palliative, G. Pain Relief Study, Alleviating the access abyss in palliative care and pain relief-an imperative of universal health coverage: the Lancet Commission report, Lancet, 391 (2018) 1391–1454.

[3] J.M. Hagel, P.J. Facchini, Biochemistry and occurrence of o-demethylation in plant metabolism, Front Physiol, 1 (2010) 14.

[4] S. Galanie, K. Thodey, I.J. Trenchard, M. Filsinger Interrante, C.D. Smolke, Complete biosynthesis of opioids in yeast, Science, 349 (2015) 1095–1100.

[5] I. Desgagne-Penix, S.C. Farrow, D. Cram, J. Nowak, P.J. Facchini, Integration of deep transcript and targeted metabolite profiles for eight cultivars of opium poppy, Plant Mol Biol, 79 (2012) 295–313.

[6] L. Pei, B.S. Wang, J. Ye, X.D. Hu, L.H. Fu, K. Li, Z.Y. Ni, Z.L. Wang, Y.J. Wei, L.Y. Shi, Y. Zhang, X. Bai, M.W. Jiang, S.H. Wang, C.L. Ma, S.J. Li, K.H. Liu, W.S. Li, B. Cong, Genome and transcriptome of Papaver somniferum Chinese landrace CHM indicates that massive genome expansion contributes to high benzylisoquinoline alkaloid biosynthesis, Hortic Res-England, 8 (2021).

[7] S. Pathak, D. Lakhwani, P. Gupta, B.K. Mishra, S. Shukla, M.H. Asif, P.K. Trivedi, Comparative Transcriptome Analysis Using High Papaverine Mutant of Papaver somniferum Reveals Pathway and Uncharacterized Steps of Papaverine Biosynthesis, Plos One, 8 (2013).

[8] U.V.T. Hong, M. Tamiru-Oli, B. Hurgobin, C.R. Okey, A.R. Abreu, M.G. Lewsey, Insights into opium poppy (Papaver spp.) genetic diversity from genotyping-by-sequencing analysis, Sci Rep, 12 (2022) 111.

[9] S.C. Farrow, J.M. Hagel, G.A. Beaudoin, D.C. Burns, P.J. Facchini, Stereochemical inversion of (S)-reticuline by a cytochrome P450 fusion in opium poppy, Nat Chem Biol, 11 (2015) 728–732.

[10] T. Winzer, M. Kern, A.J. King, T.R. Larson, R.I. Teodor, S.L. Donninger, Y. Li, A.A. Dowle, J. Cartwright, R. Bates, D. Ashford, J. Thomas, C. Walker, T.A. Bowser, I.A. Graham, Plant science. Morphinan biosynthesis in opium poppy requires a P450-oxidoreductase fusion protein, Science, 349 (2015) 309–312.

[11] R.S. Allen, A.G. Millgate, J.A. Chitty, J. Thisleton, J.A. Miller, A.J. Fist, W.L. Gerlach, P.J. Larkin, RNAi-mediated replacement of morphine with the nonnarcotic alkaloid reticuline in opium poppy, Nat Biotechnol, 22 (2004) 1559–1566.

[12] M.C. Zhong, X.D. Jiang, G.Q. Yang, W.H. Cui, Z.Q. Suo, W.J. Wang, Y.B. Sun, D. Wang, X.C. Cheng, X.M. Li, X. Dong, K.X. Tang, D.Z. Li, J.Y. Hu, Rose without prickle: genomic insights linked to moisture adaptation, Natl Sci Rev, 8 (2021).

[13] T.C. Boothby, R.S. Zipper, C.M. van der Weele, S.M. Wolniak, Removal of retained introns regulates translation in the rapidly developing gametophyte of Marsilea vestita, Dev Cell, 24 (2013) 517–529.

[14] J. Feng, W. Jia, S. Lv, H. Bao, F. Miao, X. Zhang, J. Wang, J. Li, D. Li, C. Zhu, S. Li, Y. Li, Comparative transcriptome combined with morpho-physiological analyses revealed key factors for differential cadmium accumulation in two contrasting sweet sorghum genotypes, Plant Biotechnol J, 16 (2018) 558–571.

[15] N.Q. Dong, H.X. Lin, Contribution of phenylpropanoid metabolism to plant development and plant-environment interactions, J Integr Plant Biol, 63 (2021) 180–209.

[16] X. Zhang, C.J. Liu, Multifaceted regulations of gateway enzyme phenylalanine ammonia-lyase in the biosynthesis of phenylpropanoids, Mol Plant, 8 (2015) 17–27.

[17] C.M. Fraser, C. Chapple, The phenylpropanoid pathway in Arabidopsis, Arabidopsis Book, 9 (2011) e0152.

[18] T. Vogt, Phenylpropanoid biosynthesis, Mol Plant, 3 (2010) 2–20.

[19] L. Pei, B. Wang, J. Ye, X. Hu, L. Fu, K. Li, Z. Ni, Z. Wang, Y. Wei, L. Shi, Y. Zhang, X. Bai, M. Jiang, S. Wang, C. Ma, S. Li, K. Liu, W. Li, B. Cong, Genome and transcriptome of Papaver somniferum Chinese landrace CHM indicates that massive genome expansion contributes to high benzylisoquinoline alkaloid biosynthesis, Hortic Res, 8 (2021) 5.

[20] D. Koenig, J.M. Jimenez-Gomez, S. Kimura, D. Fulop, D.H. Chitwood, L.R. Headland, R. Kumar, M.F. Covington, U.K. Devisetty, A.V. Tat, T. Tohge, A. Bolger, K. Schneeberger, S. Ossowski, C. Lanz, G. Xiong, M. Taylor-Teeples, S.M. Brady, M. Pauly, D. Weigel, B. Usadel, A.R. Fernie, J. Peng, N.R. Sinha, J.N. Maloof, Comparative transcriptomics reveals patterns of selection in domesticated and wild tomato, Proc Natl Acad Sci U S A, 110 (2013) E2655–2662.

[21] N. Chaturvedi, M. Singh, A.K. Shukla, A.K. Shasany, K. Shanker, R.K. Lal, S.P. Khanuja, Comparative analysis of Papaver somniferum genotypes having contrasting latex and alkaloid profiles, Protoplasma, 251 (2014) 857–867.

[22] K. Hori, Y. Yamada, R. Purwanto, Y. Minakuchi, A. Toyoda, H. Hirakawa, F. Sato, Mining of the Uncharacterized Cytochrome P450 Genes Involved in Alkaloid Biosynthesis in California Poppy Using a Draft Genome Sequence, Plant Cell Physiol, 59 (2018) 222–233.

[23] Q. Li, S. Ramasamy, P. Singh, J.M. Hagel, S.M. Dunemann, X. Chen, R. Chen, L. Yu, J.E. Tucker, P.J. Facchini, S. Yeaman, Gene clustering and copy number variation in alkaloid metabolic pathways of opium poppy, Nat Commun, 11 (2020) 1190.

[24] L. Yuan, E. Grotewold, Plant specialized metabolism, Plant Sci, 298 (2020) 110579.

[25] Q. Zhao, Lignification: Flexibility, Biosynthesis and Regulation, Trends Plant Sci, 21 (2016) 713–721.

[26] J. Le Roy, B. Huss, A. Creach, S. Hawkins, G. Neutelings, Glycosylation Is a Major Regulator of Phenylpropanoid Availability and Biological Activity in Plants, Frontiers in Plant Science, 7 (2016).

[27] R. Nakabayashi, K. Saito, Integrated metabolomics for abiotic stress responses in plants, Curr Opin Plant Biol, 24 (2015) 10–16.

[28] W. Dong, L. Sun, B. Jiao, P. Zhao, C. Ma, J. Gao, S. Zhou, Evaluation of aphid resistance on different rose cultivars and transcriptome analysis in response to aphid infestation, BMC Genomics, 25 (2024) 232.

[29] Z.L. Wang, S. Wang, Y. Kuang, Z.M. Hu, X. Qiao, M. Ye, A comprehensive review on phytochemistry, pharmacology, and flavonoid biosynthesis of Scutellaria baicalensis, Pharm Biol, 56 (2018) 465–484.

